# Fragmented gut–airway microbial networks and airway Moraxella clusters in preschool wheeze

**DOI:** 10.64898/2026.06.25.734489

**Authors:** Silvia Gschwendtner, Nicole Maison, Sabina Illi, Erika von Mutius, Ilona Rosenboom, Burkhard Tummler, Anna-Maria Dittrich, Markus Weckmann, Mustafa Abdo, Benjamin Waschki, Matthias V. Kopp, Gesine Hansen, Folke Brinkmann, Klaus Rabe, Bianca Schaub, Michael Schloter, the ALLIANCE Study Group

## Abstract

Early-life wheezing in children has been associated with microbial alterations along the gut-airway axis, yet studies simultaneously investigating bacterial communities in both compartments remain scarce. The aim of this cross-sectional exploratory pilot study (n=25) was to characterize and compare nasal and stool bacterial communities in preschool children aged 1–4 years with recurrent wheezing and healthy controls using 16S rRNA gene metabarcoding.

Across participants, nasal and stool bacteriomes were highly individualized and taxonomically diverse. Overall richness, evenness, and community composition did not differ significantly between healthy children and wheezers in either compartment. However, wheezers displayed markedly higher within-group variability, particularly in nasal communities. Stratification based on microbiome similarity to healthy samples revealed increased *Moraxella* and reduced commensal genera including *Prevotella* spp. and *Veillonella*, along with lower richness and evenness (all p<0.001) in nasal samples with divergent bacterial communities. Stool alterations were more subtle but included trends toward reduced *Bacteroides*, *Faecalibacterium*, and *Alistipes* in wheezers more divergent from healthy controls.

Community assembly in both compartments was largely governed by stochastic processes but accompanied by less complex and more fragmented bacterial interaction networks in wheezing children. Cross-compartment correlations were also altered, most prominently involving stool *Lactococcus* showing stronger and more numerous correlations with nasal taxa in wheezers than in healthy controls. Divergent wheezers exhibited distinct modular network structure and cross-compartment profiles, consistent with a differentiated microbial organization.

Together, these findings suggest compartment-specific differences in microbial interaction patterns across the gut–airway axis in early-life wheezing, despite limited differences in overall community diversity.

**Take home message:** Preschool wheezers showed fragmented gut–airway microbial networks and *Moraxella*-associated airway community stratification despite limited differences in overall diversity.

## Introduction

Human microbiota play a central role in regulating local and systemic immune function. A prominent example of such systemic crosstalk is the axis between the gut and the respiratory system, describing bidirectional communication between intestinal and airway microbiomes mediated by microbial metabolites and immune modulation [1–3]. Increasing evidence indicates that this axis is associated with asthma pathogenesis, particularly during early life when immune development is highly sensitive to environmental influences [4, 5].

The gut microbiome begins to establish at birth and undergoes rapid succession during infancy. In human studies, stool samples commonly serve as a non-invasive proxy for intestinal microbial communities. Early colonization is influenced by mode of delivery, diet, and environmental exposures [6]. During early infancy, *Bifidobacterium* species dominate, particularly in breastfed infants [7, 8]. With the introduction of complimentary feeding, microbial diversity increases with greater representation of *Bacteroides* and Firmicutes including *Blautia*, *Faecalibacterium*, and *Roseburia* [7]. By approximately three years of age, the microbiome stabilizes toward an adult-like configuration dominated by *Bacteroides, Prevotella,* and various Firmicutes [8, 9].

The airway microbiome develops along a similarly dynamic trajectory. Given the continuity of the respiratory tract, nasal microbiota may provide insight into early upper-airway colonization that contributes to microbial immigration into the lower airways and may influence respiratory immune development [4, 10–12]. In early infancy, nasal communities are typically dominated by *Staphylococcus* and *Corynebacterium* [13], while *Alloiococcus* and *Moraxella* increase during the first year of life [12]. Early colonization with *Streptococcus* and *Haemophilus* has been associated with viral respiratory tract infections, although these genera may also appear during normal microbial succession [12]. With increasing age, airway communities diversify further, with taxa such as *Dolosigranulum* and *Corynebacterium* becoming more prominent by two to three years of age [14]. As in the gut, early microbial composition and diversity in the airways have been linked to immune development and allergic sensitization [15].

Consistent with these observations, early microbial dysbiosis in either the intestinal or airway microbiome has been associated with wheezing illnesses and subsequent asthma development [12, 16–20]. While these findings support the concept of an axis of the gut and the respiratory system, most studies have examined the gut or respiratory microbiome separately, and investigations assessing microbial communities of both organs in parallel remain rare.

To address this gap, we characterized microbial communities of the gut and upper airways in children aged 1–4 years with recurrent wheezing (n=17) and compared them with healthy controls (n=8) in a cross-sectional exploratory pilot study. Stool samples and nasal swabs served as proxies for intestinal and upper airway microbiomes, respectively. Bacterial community composition and diversity were assessed using a metabarcoding approach, enabling parallel profiling of microbial communities in both compartments and facilitating the identification of microbial signatures associated with early-life wheezing and asthma risk.

## Materials and Methods

### Study design

The ALLIANCE cohort of the German Center for Lung Research (DZL) is a prospective multicenter asthma cohort recruiting in five pediatric specialist centers (Hannover, Luebeck, Munich, Marburg and Cologne) and two adult specialist centers (LungenClinic Grosshansdorf and Research Centre Borstel), registered at ClinicalTrials.gov (pediatric arm NCT02496468, adult arm NCT02419274) and ethically approved across all study centers. Parents of study participants aged <18 years and study participants aged >=18 years gave written informed consent.

Eligibility for this exploratory pilot study required age 1–4 years, availability of paired nasal swab and stool samples, and wheeze status. The wheezer group comprised 19 children from a total of 221 wheezers with ≥2 wheezing episodes in the preceding 12 months based on parental reports. Two wheezers were excluded due to absence of bacterial sequencing reads from nasal swabs; corresponding stool samples were also excluded to maintain paired analyses, resulting in 17 wheezers. Eight age- and sex-matched healthy controls (HC) were recruited from 54 controls if never diagnosed with asthma or preschool wheeze, but irrespective of other allergic diseases. Patient characteristics are summarized in Table 1. Details regarding study design, definitions of parameters, inclusion criteria, and methods are specified in Supplementary Material and published previously [21].

**Table 1.** Summary of baseline characteristics of the study cohort.

| Characteristic | Overall<br>N = 25 | healthy<br>N = 8 | wheeze<br>N = 17 | p value <sup>1</sup> |
| --- | --- | --- | --- | --- |
| Age (year) | 2.70 (1.90-3.30) | 2.65 (1.90-2.85) | 3.00 (1.90-3.40) | 0.209 |
| Female | 8 (32%) | 4 (50%) | 4 (24%) | 0.359 |
| BMI (kg/m<sup>3</sup>) | 16.80 (15.50-17.94) | 16.50 (15.35-17.30) | 16.98 (15.50-18.20) | 0.268 |
| Any sibling | 16 (64%) | 4 (50%) | 12 (71%) | 0.394 |
| Daycare | 23 (92%) | 8 (100%) | 15 (88%) | >0.999 |
| Pet in household (cat/dog) | 9 (36%) | 2 (25%) | 7 (41%) | 0.661 |
| Asthma or recurrent<br>spastic/obstructive bronchitis -<br>DD | 2 (8.3%) | 0 (0%) | 2 (13%) | 0.536 |
| Missing | 1 | 0 | 1 |  |
| Hayfever - DD | 1 (4.2%) | 0 (0%) | 1 (5.9%) | >0.999 |
| Missing | 1 | 1 | 0 |  |
| Eczema - DD | 2 (8.3%) | 0 (0%) | 2 (13%) | 0.536 |
| Missing | 1 | 0 | 1 |  |
| Food allergy - DD | 0 (0%) | 0 (0%) | 0 (0%) | -- <sup>2</sup> |
| Missing | 1 | 1 | 0 |  |
| Birthmode |  |  |  | 0.095 |
| Vaginal delivery | 15 (60%) | 7 (88%) | 8 (47%) |  |
| Primary cesarean section | 6 (24%) | 0 (0%) | 6 (35%) |  |
| Secondary cesarean section | 2 (8.0%) | 1 (13%) | 1 (5.9%) |  |
| Operative vaginal delivery | 2 (8.0%) | 0 (0%) | 2 (12%) |  |
| Any exclusive breastfeeding ever | 19 (76%) | 5 (63%) | 14 (82%) | 0.344 |
| Any hypoallergenic formula ever | 9 (36%) | 1 (13%) | 8 (47%) | 0.182 |
| Prior RSV infection ever | 7 (32%) | 0 (0%) | 7 (47%) | 0.051 |
| Missing | 3 | 1 | 2 |  |
| Antibiotics given for recurrent<br>cough ever | 1 (33%) | NA | 1 (33%) | -- <sup>2</sup> |
| Missing | 22 | 8 | 14 |  |
| Systemic steroids ever | 9 (53%) | NA | 9 (53%) | -- <sup>2</sup> |
| Missing | 8 | 8 | 0 |  |
| ICS ever | 13 (76%) | NA | 13 (76%) | -- <sup>2</sup> |
| Missing | 8 | 8 | 0 |  |
| Hb (g/dl) | 11.60 (11.40-12.30) | 11.60 (11.30-12.30) | 11.90 (11.45-12.25) | 0.365 |
| Missing | 2 | 1 | 1 |  |
| Thrombocytes (10<sup>3</sup>/μl) | 374 (328-443) | 328 (263-367) | 403 (372-443) | 0.045 |
| Missing | 2 | 1 | 1 |  |
| Leucocytes (10<sup>3</sup>/μl) | 9.44 (8.23-11.70) | 10.00 (5.24-14.00) | 9.27 (8.24-11.65) | 0.841 |
| Missing | 2 | 1 | 1 |  |
| Neutrophils (%) | 40 (33-55) | 38 (31-58) | 42 (36-55) | 0.947 |
| Missing | 2 | 1 | 1 |  |
| Lymphocytes (%) | 42 (34-54) | 53 (27-60) | 40 (36-47) | 0.593 |
| Missing | 2 | 1 | 1 |  |
| Eosinophils (%) | 1.95 (1.00-3.00) | 2.50 (1.00-3.50) | 1.50 (0.78-2.40) | 0.243 |
| Missing | 3 | 1 | 2 |  |
| Basophils (%) | 0.20 (0.00-0.70) | 0.00 (0.00-0.20) | 0.55 (0.08-1.00) | 0.036 |
| Missing | 4 | 1 | 3 |  |
| IgE (IU/ml) | 39.0 (9.1-73.8) | 47.5 (24.0-61.4) | 19.2 (9.0-82.0) | 0.583 |
| Missing | 2 | 0 | 2 |  |
| Atopy (spec. IgE to any allergen<br>≥0.7 kU/L) | 8 (32%) | 3 (38%) | 5 (29%) | >0.999 |

Values are summarized as median and interquartile range (Q1-Q3) in brackets, or as total numbers per condition level and percentage in brackets.
DD = parental report of doctor's diagnosis
<sup>1</sup>Kruskal-Wallis rank sum test; Fisher's exact test
<sup>2</sup>p value not calculated due to insufficient variability (single-level data) and/or missing values

### Microbiome analysis

Details on microbiome sample processing and subsequent data analysis are provided in Supplementary Material. Briefly, DNA extraction was performed via ultrasonication (nose) and a phenol-chloroform-based protocol (stool), alongside with blank samples to identify potential contaminants. Amplicon sequencing of the V4 hypervariable region of the 16S rRNA gene was performed on a MiSeq Illumina instrument using the universal eubacterial primers 515F/806R [22, 23]. Sequence data generated in this study are available upon reasonable request from the authors.

Sequence processing was performed using DADA2 v 1.30 [24], followed by taxonomic assignment via SILVA v138.1. Reads classified as mitochondrial or chloroplast in origin, lacking phylum-level annotation, or detected in blank samples were excluded, resulting in a total of 3,744,773 reads assigned to 2,177 amplicon sequence variants (ASV).

All plots and statistics were performed in R version 4.5.1 (https://www.R-project.org). Subsequent bioinformatic analyses were performed to characterize microbial communities in HC and wheezers, including assessments of alpha and beta diversity, core microbiome structure, within-group variability, and microbial biomarkers (ANCOM-BC2). Additional analyses explored associations between microbial taxa, individual variability and host factors, cross-compartment correlations, microbial community assembly processes (f1NTI and RCbray framework), and bacterial co-occurrence networks (NetCoMi). Furthermore, wheezers were stratified into healthy-like and divergent microbiome profiles and their associated features were examined. Detailed protocols are provided in Supplementary Material.

## Results

### Cohort Characteristics

The study comprised 25 participants at preschool age (17 children with recurrent wheeze and 8 healthy controls (HC)), with baseline data summarized in Table 1. Demographic characteristics (age, gender), early-life and environmental exposures (e.g. mode of delivery, breastfeeding, siblings, daycare, pet contact) and most clinical history variables (allergic and respiratory conditions, prior RSV infection) did not differ between HC and wheezers, while use of antibiotics or steroids (systemic and inhaled) was only prevalent in wheezers. Wheezers exhibited higher thrombocyte and blood basophil counts, while no differences were observed for other hematological or immunological parameters.

### Nasal and stool bacteriomes show similar bacterial diversity between groups but higher compositional variability in wheezers

Across participants, nasal and stool bacteriomes were taxonomically diverse but highly individualized. The nasal bacteriome comprised 936 ASV and stool samples 1033 ASV, with overlap but more wheezer-specific variants (Fig. 1a). Group-specific core ASV decreased rapidly with increasing prevalence thresholds in both compartments, indicating limited overlap between individuals (Fig. 1a; Supplementary Table S1). Nevertheless, alpha diversity metrics indicated comparable richness and evenness between HC and wheezers in both compartments (Supplementary Fig. S1a), and NMDS analysis of weighted UniFrac distances showed no group separation (Supplementary Fig. S1b).

**Figure 1.**
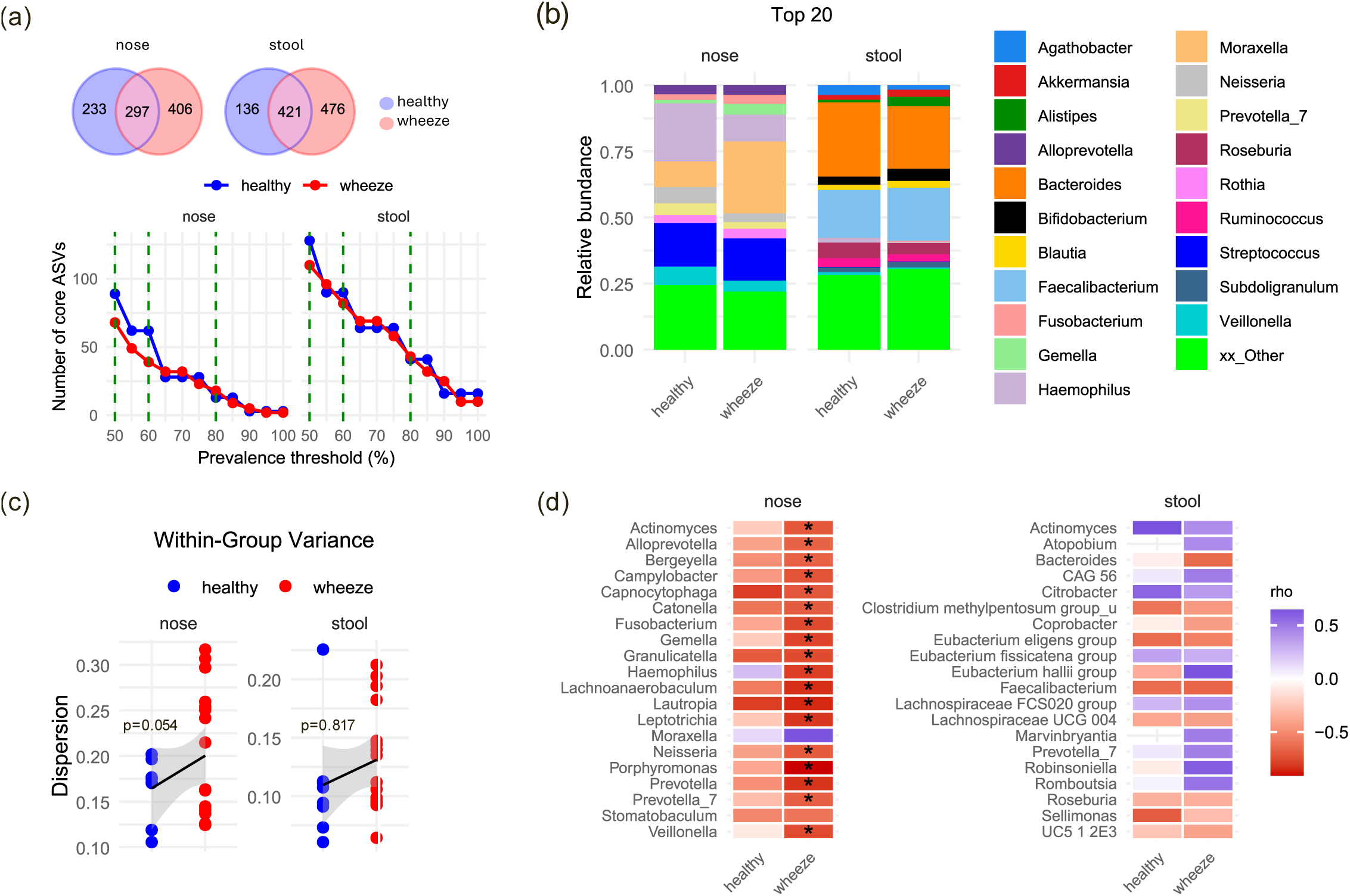
Nasal and gut microbiome composition in healthy (n=8) and wheezing (n=17) children. (a) Shared and unique ASV and core ASV across prevalence thresholds. Top: Venn diagrams showing shared and group-specific ASV in nasal and stool samples. Bottom: Number of core ASV across prevalence thresholds (50–100%), defined as ASV present in ≥X% of samples within each study group. (b) Relative abundance of the 20 most abundant genera. No significant differences in genus abundance were detected between groups (ANCOM-BC2; Benjamini–Hochberg-adjusted p<0.05). (c) Within-group variation (beta-dispersion) calculated from weighted UniFrac distances. Statistical significance was assessed using Levene’s test and confirmed using generalized linear mixed-effects models with Benjamini–Hochberg adjustment for multiple testing. (d) Spearman correlations between beta-dispersion and bacterial genera. Only the 20 strongest correlations, ranked by Spearman’s correlation coefficient (rho), are shown. Asterisks indicate statistically significant correlations (Benjamini–Hochberg-adjusted p<0.05). Correlations were calculated using Spearman’s rank correlation test.

Nasal bacterial communities were dominated by *Haemophilus*, *Moraxella*, *Streptococcus*, *Veillonella*, and *Prevotella*-related taxa (Fig. 1b). *Moraxella* tended to be more abundant in wheezers (p>0.05) and occasionally dominated individual profiles (Supplementary Fig. S2). Among 11 *Moraxella* ASV, one variant closely related to *Moraxella nonliquefaciens* predominated across samples (Supplementary Fig. S3) but was more consistently present in HC despite lower relative abundance (Supplementary Table S1). Beta-dispersion indicated greater within-group variability in the nasal bacteriome of wheezers compared with HC (p=0.054; Fig. 1c) and subgroup-specific associations with nasal genera. In wheezers, *Neisseria, Prevotella* spp., and *Veillonella* were negatively correlated, while *Moraxella* showed a non-significant positive association (Fig. 1d). Comparable non-significant trends were seen in controls.

Stool communities showed higher richness (p<0.001) and evenness (p=0.004) than nasal samples (Fig. S1) and were dominated by *Bacteroides* and *Faecalibacterium* (>43% of reads; Fig. 1b). Differences between groups were modest, although several commensals like *Roseburia* tended to be reduced in wheezers. Stool within-group dispersion was lower compared with nasal samples (p<0.001) but similarly tended to be higher in wheezers (Fig. 1c). This variability was associated with higher *Romboutsia* and *Prevotella_7* and lower *Bacteroides*, *Faecalibacterium*, and *Roseburia*, although not significant after correction for multiple comparisons (Fig. 1d).

Despite substantial inter-individual variability, several host-microbiome associations were observed. In the nose, leukocyte levels correlated negatively with *Corynebacterium, Dolosigranulum* and *Moraxella* (Supplementary Fig. S4a). In stool, immune parameters were not associated with bacterial diversity. However, age correlated negatively with *Bifidobacterium*, while *Ruminococcus* and *Agathobacter* were associated with gender. Beta-dispersion, used as a measure of individual variability, was not related with measured host parameters (Supplementary Fig. S4b).

### Microbial community divergence identifies distinct wheezer subgroups with divergent nasal communities enriched in Moraxella

To explore heterogeneity among wheezers, samples were stratified into healthy-like and community-divergent based on the Euclidean distance of each sample’s bacterial community to the HC centroid in ordination space (90% quantile threshold). This classification reflects similarity in community structure and was not associated with differences in demographic or clinical parameters (Supplementary Table S2).

In the nasal bacteriome, 9 of 17 wheezers showed a healthy-like bacterial profile, whereas 8 were classified as divergent. Divergent wheezers displayed higher individual variability but lower alpha diversity measures (observed ASV, Shannon index, evenness) (Fig. 2a, Supplementary Table S3), increased *Moraxella* abundance, and reduced abundance of commensals such as *Alloprevotella*, *Prevotella_7*, and *Veillonella* at both genus (Fig. 2b) and ASV level (Fig. 2c).

**Figure 2.**
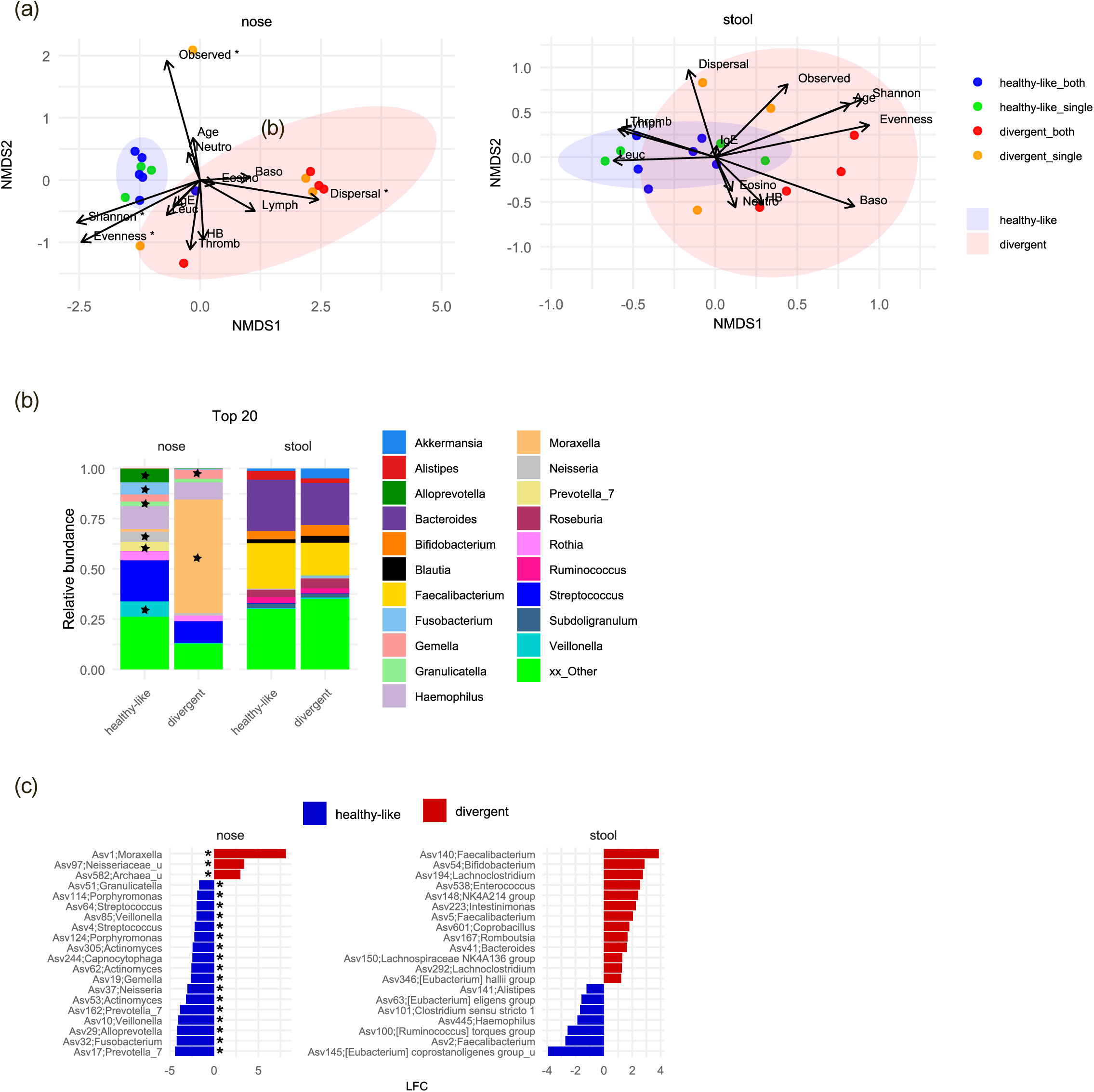
Microbial signatures of healthy-like and community-divergent wheezers. (a) Non-metric multidimensional scaling (NMDS) ordination based on Bray–Curtis distances, with alpha-diversity metrics (observed ASV, evenness and Shannon index), beta-dispersion and immune variables fitted as environmental vectors. Asterisks indicate vectors significantly associated with community composition (permutation test within the envfit function, p<0.05). Vector length reflects the strength of the association with the ordination, and vector direction indicates the gradient of increasing values across the NMDS space. Blue and red symbols represent subjects classified as healthy-like and community-divergent wheezers, respectively, in both nasal and gut microbiomes; green and orange symbols represent subjects classified as healthy-like and community-divergent wheezers, respectively, in only one compartment (nasal or gut). (b) Relative abundance of the 20 most abundant genera. Asterisks indicate significant differences between healthy-like and community-divergent wheezers (ANCOM-BC2; Benjamini–Hochberg-adjusted p<0.05). (c) Log-fold changes (LFC) estimated by ANCOM-BC2 for the 20 ASV with the largest effect sizes differentiating healthy-like and community-divergent wheezers. Asterisks indicate significant differences after Benjamini–Hochberg adjustment (adjusted p<0.05).

In stool bacterial communities, 7 wheezers were classified as divergent based on distance from the healthy centroid. Differences between subgroups were less pronounced than in the nose, but divergent individuals tended to show reduced *Bacteroides*, *Faecalibacterium*, and *Alistipes* (Fig. 2b). As in nasal communities, baseline parameters did not differ between stool-based subgroups (Supplementary Table S2).

Partial overlap between compartments was observed: four of the eight nasal-divergent wheezers were also classified as divergent in stool, whereas the remaining individuals showed healthy-like stool profiles. Similarly, six of nine healthy-like wheezers displayed concordant bacterial signatures in both compartments.

### Stochastic assembly dominates nasal and stool bacteriomes, while wheezing is associated with fragmented bacterial interaction networks

To investigate processes shaping nasal and stool bacterial communities, we quantified the relative influence of stochastic and deterministic assembly mechanisms using βNTI and RCbray metrics (Fig. 3). Stochastic processes dominated bacterial community assembly in both compartments. In HC, nasal communities were primarily shaped by dispersal limitation (restricted microbial exchange between individuals/environments) and drift (random fluctuations in microbial abundances). Wheezers additionally exhibited deterministic processes (variable selection), suggesting host- or environment-related selection of specific microbial taxa. Stool communities were also mainly structured by dispersal limitation, with selective processes playing only a minor role in HC but gained importance in wheezers.

**Figure 3.**
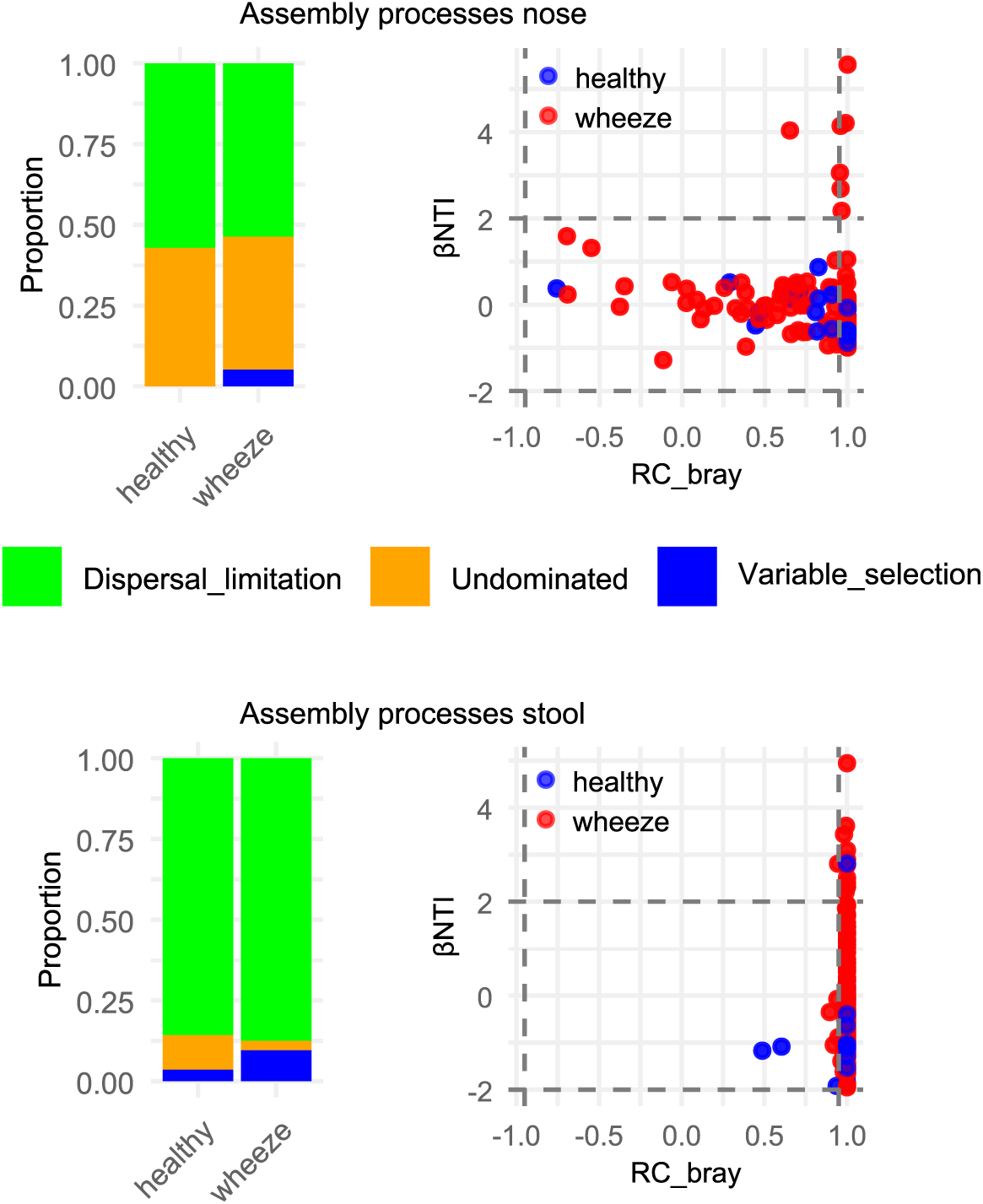
Community assembly processes in the nasal and gut microbiomes of healthy (n=8) and wheezing (n=17) children. Community assembly mechanisms were inferred using β-nearest taxon index (βNTI) and Raup–Crick Bray–Curtis (RCbray) metrics. Thresholds of |βNTI|=2 and |RCbrayl=0.95 are indicated by horizontal and vertical dashed lines, respectively.

Co-occurrence network analysis revealed pronounced differences between HC and wheezers, with similar trends in nasal and stool bacteriomes (Fig. 4). Wheezing was associated with reduced network complexity in both compartments, reflected by fewer nodes, modules, and lower modularity. In contrast, clustering coefficient and centrality measures (degree, betweenness, closeness) were increased. Compartment-specific differences in path length, and proportion of positive edges were observed but less pronounced than the overall pattern of network simplification and reorganization. Notably, the divergent subgroup showed higher modularity and lower clustering and centrality measures than the overall wheezer group,consistent with a distinct microbial configuration. In the nose, network hubs shifted from ASV assigned to commensals like *Staphylococcus* and *Cutibacterium* in HC to *Leptotrichia*, *Fusobacterium*, and *Prevotella* spp. in wheezers. A similar trend was observed in stool, with hubs transitioned from *Veillonella* and *Roseburia* toward Lachnospiraceae and *Ruminococcus*.

**Figure 4.**
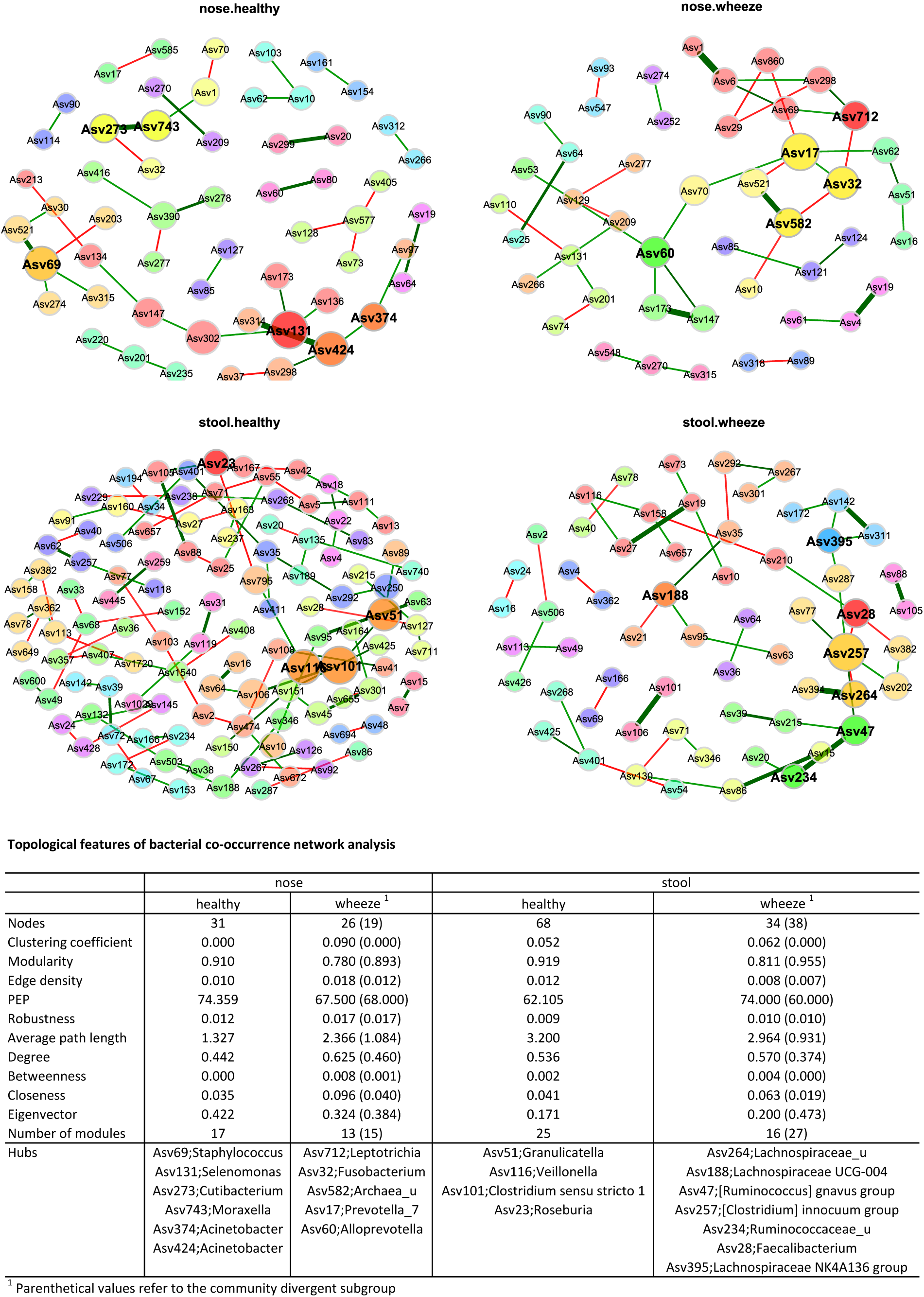
Bacterial interaction networks in healthy (n=8) and wheezing (n=17) children. Bacterial co-occurrence networks were constructed from core ASV, defined as ASV present in ≥50% of subjects, in nasal and stool samples from healthy and wheezing children. Nodes represent ASV and edges represent inferred associations between pairs of ASV estimated using SpiecEasi (green: positive associations; red: negative associations). Modules, shown in different colours, represent groups of highly interconnected nodes. Hub taxa are highlighted in bold, and node size is proportional to eigenvector centrality. The accompanying table summarises key network topological characteristics, including the number of nodes, clustering coefficient, modularity, positive edge percentage (PEP), edge density, robustness, average path length, centrality measures (degree, betweenness, closeness and eigenvector centrality), and the number of modules.

### Altered cross-compartment correlations between nasal and stool bacterial communities in wheezers

Spearman correlation analysis revealed distinct nasal–stool co-variation patterns between HC and wheezers (Fig. 5), with 42 significant correlations in HC (lρl>=0.97) and 53 in wheezers (lρl>=0.82), mostly positive.

**Figure 5.**
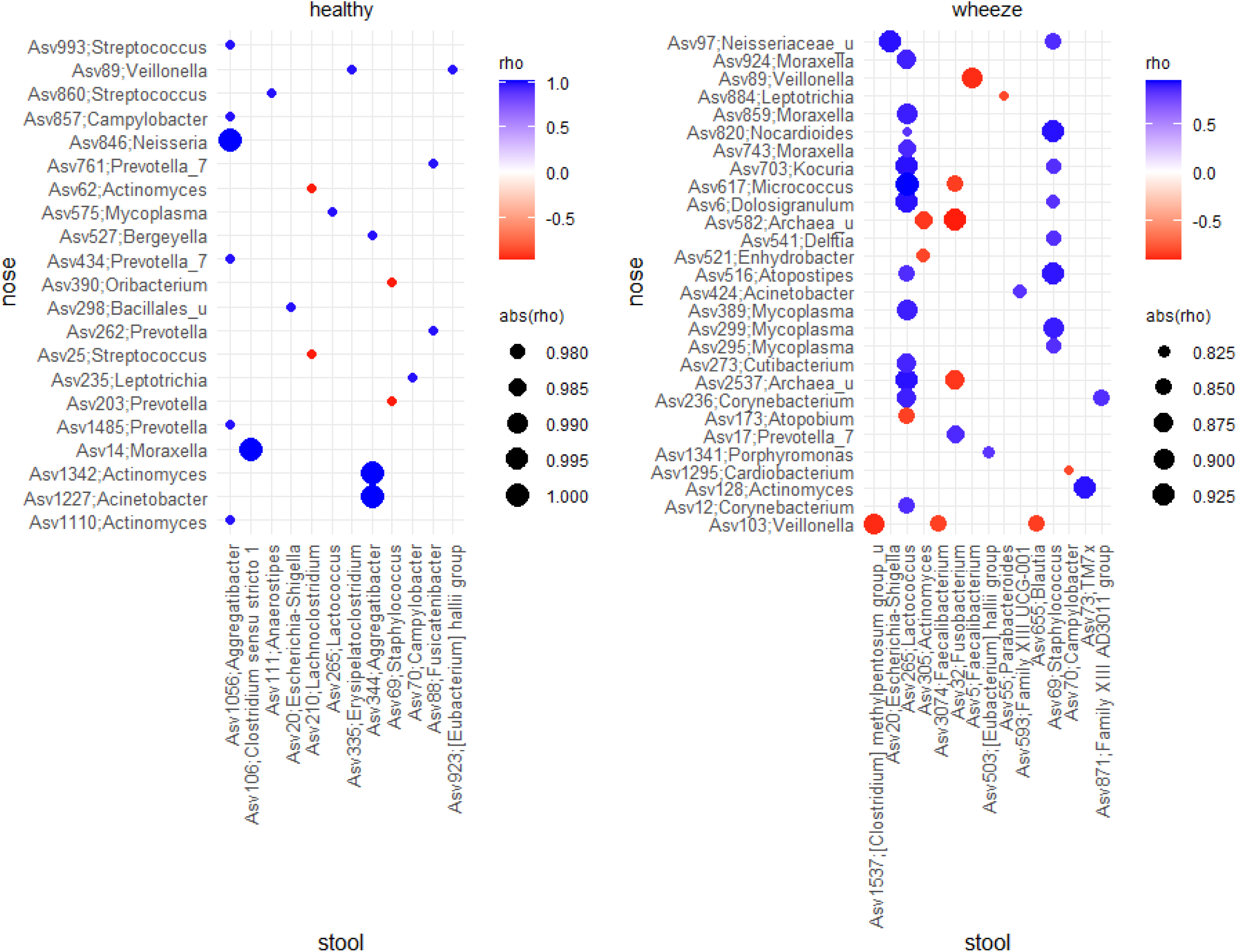
Pairwise cross-compartment correlations in healthy (n=8) and wheezing (n=17) children. Spearman correlations were calculated between individual ASV in nasal and stool samples. Only statistically significant correlations are shown (Benjamini–Hochberg-adjusted p<0.05).

In HC, positive correlations were mainly observed among commensals, including nasal ASV assigned to *Prevotella*, *Neisseria*, *Actinomyces*, and *Veillonella* and stool ASV related to *Aggregatibacter*. In addition, stool ASV210 (*Lachnoclostridium)* and ASV69 (*Staphylococcus*) were negatively associated with nasal ASV assigned to *Actinomyces, Streptococcus*, and *Prevotella*, suggesting coordinated variation across compartments in HC.

In wheezers, cross-compartment associations differed markedly. Several nasal ASV assigned to *Corynebacterium*, *Dolosigranulum*, and *Moraxella* showed strong positive correlations with stool *Lactococcus* ASV265 (ρ=0.82–0.94). Additional associations included positive correlations between nasal *Dolosigranulum* and *Mycoplasma* ASV and stool *Staphylococcus* (ASV69), as well as negative correlations between nasal *Veillonella* and stool *Faecalibacterium* and *Blautia.* Furthermore, stool *Escherichia–Shigella* (ASV20) was positively associated with nasal *Neisseriaceae_u (ASV97)*. These correlations were largely absent in the divergent subgroup, showing mainly associations between stool *Escherichia-Shigella*, *Faecalibacterium,*, *Rothia* and various nasal taxa (Supplementary Fig. S4).

## Discussion

### Nasal bacterial diversity in wheezers is characterized by increased variability and Moraxella-associated community shifts

In this cross-sectional study, we analysed nasal and stool bacterial communities in preschool children with recurrent wheezing and HC across the gut–airway axis. While overall bacterial richness and community composition were comparable between groups, wheezers showed higher inter-individual variability in nasal bacterial communities.

Moraxella, a common colonizer of the upper airways, showed a strong but not significant correlation with nasal community patterns. Previous studies have reported contrasting roles of *Moraxella* in respiratory health. Some studies describe it as a typical component of healthy airway communities [13, 25], whereas others link its predominance to respiratory tract infections and wheezing episodes [12, 26]. Our results suggest that the ecological context of *Moraxella* is more important than its mere presence, In HC it occurred at low abundance within diverse communities, whereas wheezers often showed *Moraxella*-dominated profiles associated with reduced richness and evenness. Similar cluster-like community patterns have previously been described in airway microbiota during early-life [26, 27] and at school age [28]. Comparable alterations are seen in chronic airway diseases such as cystic fibrosis and bronchiectasis, where reduced diversity is accompanied by dominance of opportunistic pathogens including *Pseudomonas aeruginosa*, and *Haemophilus influenzae*, and is associated with more severe disease [29, 30]. These observations suggest that *Moraxella* may act as an opportunistic taxon that expands when microbial community balance is disturbed, potentially following viral infections or changes in local immune responses. Future studies should stratify children by prior infection status and integrate multi-omics data and immunological profiles, to better disentangle host–microbiome interactions driving *Moraxella* expansion.

### Stool bacterial diversity remains comparatively stable beyond infancy

In contrast to nose, stool bacterial communities showed only modest alterations between wheezing and HC, consistent with previous evidence that intestinal microbial signatures associated with asthma risk are strongest in early infancy rather than later childhood [17, 19, 20]. Longitudinal studies have demonstrated that intestinal bacterial communities often normalize during the first year of life [16, 18], which may explain why differences in preschool-aged children are less pronounced.

Our data are consistent with the concept of a developmental window during early infancy in which intestinal bacterial colonization has the strongest influence on immune maturation and asthma susceptibility. By preschool age, dietary transitions and environmental exposures may contribute to convergence of intestinal bacterial communities between wheezers and HC. However, antibiotic exposure, a known determinant of gut microbial diversity and maturation in early life, may also act as a potential cofactor influencing these patterns [31, 32]. At the same time, the relatively small cohort size and cross-sectional design limit the ability to detect subtle associations between stool bacterial diversity and wheezing phenotypes. Larger longitudinal cohorts are required to determine whether intestinal microbial signatures associated with early-life wheezing persist or resolve during later childhood, and how compartment-specific microbial patterns relate to asthma development, systemic metabolomic and immune alterations, and their potential reversibility through targeted interventions.

### Ecological restructuring of bacterial communities in wheezers

Beyond taxonomic composition, our ecological analyses revealed altered community organization between HC and wheezers. In both compartments, community assembly was dominated by stochastic processes, consistent with ecological models describing microbial colonization as a balance between dispersal-driven processes and host-mediated selection [33]. However, nasal and stool communities in wheezers showed increased deterministic contribution, potentially reflecting host-driven selection such as airway inflammation, viral infections, or local immune responses [27].

Network analysis further indicated reduced modularity and increased fragmentation of bacterial interaction networks in wheezers, reflecting a reduction in overall network complexity. Such simplification has previously been associated with disturbed microbial ecosystems and lower resilience to environmental perturbations [34, 35]. Cooperative microbial interactions, including metabolic cross-feeding and signalling processes, are thought to stabilize host-associated microbial communities [36, 37]. Their reduction may therefore contribute to microbial ecosystems that are more susceptible to instability or opportunistic expansion of individual taxa. Consistent with this, increased centrality measures indicate that fewer taxa occupied more influential positions, indicating a shift toward greater centralization and uneven distribution of connectivity among the remaining community members Notably, stratification of wheezers into subgroups revealed distinct modular network structure, suggesting differences in microbial community organization rather than a simple intensification of wheeze-associated dysbiosis.

Finally, cross-compartment correlations between nasal and stool taxa differed between HC and wheezers, suggesting group-specific altered gut-airway covariation. These correlations may reflect shared host or environmental influences rather than idirect microbial exchange across compartments and are consistent with the concept of the axis between the gut and the respiratory system, in which intestinal microbial metabolites such as short-chain fatty acids influence airway immune responses [38, 39]. Nevertheless, the functional mechanisms underlying these associations cannot be addressed by 16S rRNA metabarcoding alone and will require integrative approaches combining metagenomics, metabolomics, and immune profiling.

This study has several limitations that should be acknowledged. These include potential selection bias, cross-sectional design without longitudinal follow-up, and methodological constraints. The small sample size and exploratory pilot design limit statistical power and the generalizability of the findings. Its cross-sectional nature further precludes any conclusions regarding temporal dynamics or causality, and the use of convenience sampling may introduce additional selection bias. While 16S rRNA gene amplicon sequencing provides useful insights into community composition, it offers limited taxonomic resolution and does not capture strain-level variation or functional potential. Future studies in larger longitudinal cohorts, ideally incorporating metagenomics, metabolomics, and immune profiling, will be important to validate and extend these observations.

## Conclusions

Taken together, our results indicate that preschool wheezing is associated with increased variability and taxon-level restructuring of nasal bacterial communities, particularly involving *Moraxella*, while stool bacterial diversity remains stable. Altered microbial networks and cross-compartment correlations might further suggest coordinated ecological changes across mucosal sites. These findings highlight the importance of microbial community structure and ecological interactions, rather than diversity alone, when investigating microbial contributions to early-life wheezing and asthma susceptibility. Longitudinal functional microbiome studies will be necessary to clarify whether these signatures contribute to persistent wheezing or asthma development.

## Acknowledgements

The authors would like to thank Susanne Walch from the Research Unit Comparative Microbiome Analysis for her assistance with sequencing. We are thankful to our patients and healthy participants for their invaluable contribution to this work. We are indebted to our study participants and their families for participating in the study and the staff of the participating hospitals and primary care practices and our cooperation partners within the DZL for support and recruitment. We also thank the study nurses, the data managers, and all involved lab technicians.

## Data availability

The data cannot be made publicly available upon publication because they contain sensitive personal information. The data supporting the findings of this study are available upon reasonable requests from the authors.

## Ethics Statement

The ALLIANCE cohort of the German Center for Lung Research (DZL) was approved by all local ethics committees of the involved centers and registered at ClinicalTrials.gov (pediatric arm NCT02496468, adult arm NCT02419274). Parents of study participants aged <18 years and study participants aged >=18 years gave written informed consent.

## Author contributions

S. Gschwendtner and M. Schloter contributed to study conceptualization, performed data analysis, and drafted the manuscript. N. Maison, S. Illi, I. Rosenboom, B. Tummler, and M. Weckmann contributed to the methodology and provided scientific input, including critical review and revision of the manuscript. E. von Mutius, B. Tummler, G. Hansen, M. Kopp, and A.-M. Dittrich contributed to study conceptualization and provided scientific guidance, including critical review and revision of the manuscript. M. Abdo critically reviewed the manuscript. G. Hansen, F. Brinkmann, B. Schaub, K. Rabe, E. von Mutius, B. Waschki, and M. V. Kopp are principal investigators of the DZL ALL Age Asthma Cohort; they secured funding and contributed to the conception and development of the overarching research goals and aims.

## Supplementary Material

## Supplementary Methods

### Study design and procedures

The study design and procedures have been described in detail previously [1, 2]. For easier access we included this information here:

The ALLIANCE cohort is recruited at five pediatric centers (Cologne, Hannover, Lubeck, Marburg, Munich) and two adult centers (LungenClinic Grosshansdorf, Research Centre Borstel), all affiliated with the German Center for Lung Research (DZL). Recruitment started in 2013. Participants with preschool wheeze and asthma had annual study visits while healthy controls were assessed only once. Visits were postponed in case of upper respiratory tract infections, asthma exacerbations (adults) or fever >38.5°C in the past two weeks (children).

Caregivers and adult participants completed questionnaires covering demographic factors, early-life exposures, respiratory symptoms, medical history and medication use. Recorded variables included age, sex, BMI, mode of delivery, breastfeeding, siblings, pet contact (cat/dog), childcare, wheeze and cough during the previous 12 months, previous RSV infection, physician-diagnosed asthma, hay fever, eczema or food allergy, and use of hypoallergenic formula, antibiotics for recurrent cough, or inhaled/systemic corticosteroids. Definitions of clinical and lifestyle variables have been described previously [1]. Questionnaire-based outcomes were defined as follows: wheeze episode as salbutamol use due to wheeze on >2 of 7 days; exacerbation as a wheeze episode requiring systemic corticosteroids or hospitalization; and asthma control according to GINA guidelines.

Differential blood counts and total IgE were measured in routine on-site hospital laboratories. Allergen-specific IgE was assessed centrally using Euroline™ (Euroimmun, Germany) against a panel of common aeroallergens and food allergens [2]. Atopy was defined as ≥1 allergen-specific IgE ≥0.7 kU/L, based on previous findings from the ALLIANCE cohort [3]. Elevated blood eosinophil counts were defined as ≥470 cells/µl, corresponding to the 90th percentile of healthy controls in the ALLIANCE cohort.

### DNA extraction and library preparation

DNA of nasal swabs was extracted via freezing-heating cycles (5 min, 3 times) following ultrasonication (S220 Focused-ultrasonicator Covaris) with 6 cycles of 5 s at peak power=200/duty factor=2, 30 s peak power=275/duty factor=5 at 6°C [4]. DNA of stool samples was extracted from 0.4 g feces material using the phenol-chloroform-based protocol described by Lueders et al. [5] and the Precellys24 Instrument (PeqLab, Erlangen, Germany).

Amplicon sequencing of the V4 hypervariable region of the 16S rRNA gene was performed on a MiSeq Illumina instrument (MiSeq Reagent Kit v3 (600 Cycle); Illumina, San Diego, CA, USA) using the universal eubacterial primers 515F [6] and 806R [7]. PCR was done using NEBNext high fidelity polymerase (New England Biolabs, Ipswich, USA) in a total volume of 25 µl (10 ng DNA template (nose) / 5 ng DNA template (stool), 12.5 µl polymerase, 5 pmol of each primer, 2.5 µl 3% BSA) and the following PCR conditions: 5 min at 98°C; 30 cycles (nose) / 25 cycles (stool) of 10 s at 98°C, 30 s at 55 °C, 30 s at 72 °C; 5 min 72 °C. To identify potential contaminants deriving from DNA extraction and library preparation, both extraction and PCR no template control samples were processed (each 4). PCR products were purified using MagSi NGSprep Plus beads (Steinbrenner, Wiesenbach, Germany) and quantified via PicoGreen assay. Subsequently, indexing PCR was performed using the Nextera XT Index Kit v2 (Illumina, Inc. San Diego, CA, US) in a total volume of 25 µl (10 ng DNA template, 12.5 µl NEBNext high fidelity polymerase, 2.5 µl of each indexing primer) and the following PCR conditions: 30s at 98 °C; 8 cycles of 10 s at 98 °C, 30s at 55 °C, 30s at 72 °C; 5min 72 °C. Indexing PCR products were purified using MagSi NGSprep Plus beads, qualified and quantified via a Fragment Analyzer™ instrument (Advanced Analytical Technologies, Inc., Ankeny, USA) and pooled in an equimolar ratio of 4nM.

Sequence data generated in this study are available upon reasonable request from the authors.

### Sequence processing

FASTQ files were trimmed with a minimum read length of 50 using Cutadapt [8] and quality control was performed via FastQC [9]. For subsequent data analysis, the DADA2 pipeline v 1.30 [10] was used with the following trimming and filtering parameters: 20 bp were removed n-terminally and reads were truncated at position 240 (forward) and 200 (reverse), respectively, with expected error of 3 (forward) and 4 (reverse). Taxonomic analysis was performed using SILVA v138.1. Reads were excluded if classified as mitochondria or chloroplast or if the phylum was missing. All blank samples were analyzed together with biological samples but received only 0-4 sequencing reads, thus all sample reads were kept for subsequent analysis, resulting in a total amount of 3,744,773 reads (corresponding to an average of 72,014 reads per sample) assigned to 2,177 amplicon sequence variants (ASV).

### Statistical analysis

All plots and statistics were performed in R version 4.5.1 (https://www.R-project.org). Prior to the main analysis, potential covariates (age, gender, cough antibiotics, siblings, daycare, and pet contact) were tested individually for association with microbiome outcomes due to the modest sample size (n = 25). Covariates improving model fit, as judged by metrics like delta AIC, beta coefficients and CI for mixed-effect models or explained variance for PERMANOVA, were included in the final analysis.

Alpha diversity was calculated using species richness, evenness as well as Shannon diversity index. Associations with covariates were assessed using generalized linear mixed-effects models (R package glmmTMB), including study subjects as a random effect. Covariates improving model fit in pre-tests (age and gender) were retained. Analyses were performed on TMM-normalized data [11] and raw counts, with log-transformed sequencing depth as covariate for raw counts when it was significantly correlated with the response variable (assessed using cor.test). Beta diversity was analyzed using unweighted and weighted UniFrac distances on raw counts, as well as Aitchison distance on CLR-transformed data. Associations with covariates were tested via PERMANOVA, with permutations restricted within study subjects using the strata option (R package vegan). Based on pre-tests, age and gender was included as covariate. For both alpha and beta diversity analysis, p value adjustment for multiple comparisons was performed with Benjamini-Hochberg correction.

To identify microbial taxa differing between healthy and wheezing children, ANCOM-BC2 (Analysis of Compositions of Microbiomes with Bias Correction 2) was used [12], with age and gender as covariates. Phylogenetic trees for the ASV of interest were constructed including SILVA-curated reference sequences (https://www.arb-silva.de/). Sequences were aligned with ClustalW following tree optimization via the Maximum Likelihood method (GTR model) and 1000 bootstrap iterations for reliability (R packages ape, phangorn, msa).

For analysis of within-group variation, the average weighted UniFrac distance between each subject and the group centroid was calculated (beta-dispersion). Statistical analysis was performed like for alpha diversity via generalized linear mixed-effects models (R package glmmTMB) and Benjamini-Hochberg p value correction for multiple comparisons, including study subjects as a random effect and age and gender as covariates. In addition, Levene’s Test was used to test for equality of variances within the groups.

To search for associations between individual variability (beta-dispersion), demographic/lifestyle variables, hematological immune markers (hemoglobin, thrombo-, leuko-, lymphocytes, neutro-, eosino-, basophils, immunoglobulin E), and microbial taxa, correlation analyses were performed, with Pearson correlations for numerical variables and Spearman correlations for the comparison of numeric and microbiome data after CLR-transformation. P value adjustment for multiple comparisons was performed with Benjamini-Hochberg correction.

The core microbiome per group was defined as ASV or genera present in at least 80 % of subjects without setting an additional abundance cutoff.

To explore heterogeneity among wheezing children, samples were classified as healthy-like or community-divergent based on their microbiome’s distance to the centroid of healthy samples. First, Bray-Curtis distances were calculated on TSS-normalized data, and then Euclidean distances were computed from each sample to the healthy centroid in the ordination space. Wheezing samples within the 90th percentile of the healthy centroid distances were considered healthy-like, while those exceeding this threshold were classified as community-divergent. This resulted in body site-specific subgroup sizes of 9 and 8 for nasal samples and 10 and 7 for stool samples (healthy-like and community-divergent, respectively). Selected variables were fitted to the ordination space as vectors using the envfit function (R package vegan), with significance assessed via permutation tests. Further statistical analysis between those subgroups were performed as described above using generalized linear mixed-effects models (R package glmmTMB), ANCOM-BC2 and correlation analyses.

Microbial community assembly was assessed using the β-nearest taxon index (βNTI) and Raup-Crick–based Bray-Curtis (RCbray) approach following Stegen et al. [13]. Although βNTI was originally developed for macroecological systems, it is widely applied in microbial ecology because it quantifies phylogenetic turnover relative to null expectations, enabling inference of deterministic versus stochastic processes. Phylogenetic relatedness serves as a proxy for ecological similarity due to trait conservation in microbes. This framework was applied to evaluate how deterministic and stochastic processes shape microbial communities across healthy and wheezing children, where wheezing may impose strong selective pressures. For βNTI, thresholds of ±2 were used to define deterministic selection, further categorized as heterogeneous/variable selection (βNTI > +2; divergent community configurations) and homogeneous selection (βNTI < −2; convergence toward similar composition). For pairwise comparisons with lβNTIl < 2, RCbray distinguished stochastic processes: RCbray > +0.95 indicated dispersal limitation, RCbray < −0.95 indicated homogenizing dispersal, and |RCbray| ≤ 0.95 indicated undominated processes. Analyses were performed in R using the *picante* and *vegan* packages with null distributions generated from 999 randomizations. A phylogenetic tree was constructed prior to these analyses by aligning sequences with MAFFT (FFT-NS algorithm) and building the tree with FastTree under the GTR+CAT substitution model. The *ape* package was used to handle phylogenetic distances via a *phyloseq* object.

Microbial co-occurrence networks were constructed using the R package NetCoMi v1.1.0 [14], applying the “signed” transformation to convert estimated associations into dissimilarities and including only ASV present in ≥50% of samples per group. Networks were inferred with the SpiecEasi framework on CLR-transformed data, with zero replacement via multiplicative imputation and the Meinshausen-BOhlmann neighborhood selection method for network inference. Network topology was characterized by modular structure (fast-greedy clustering) and identification of hub taxa based on degree and eigenvector centrality (top 10%). Centrality measures (degree, betweenness, closeness, eigenvector) were normalized, and weighted node degrees were calculated to assess connectivity patterns across microbial communities.

To evaluate cross-compartment associations between nasal and stool microbiomes, raw counts were transformed to relative abundances and CLR-transformed. Spearman correlations between all nose-stool pairs were calculated using the rcorr function (R package Hmisc), with p-values adjusted via Benjamini-Hochberg. Significant correlations (p < 0.05) were visualized as a bipartite network (R packages igraph, ggraph), with nodes representing taxa and edges indicating significant associations. If not stated otherwise, all plots were created in R using ggplot2 [15], ggpubr [16], and eulerr [17].

---

**Figure S1.**
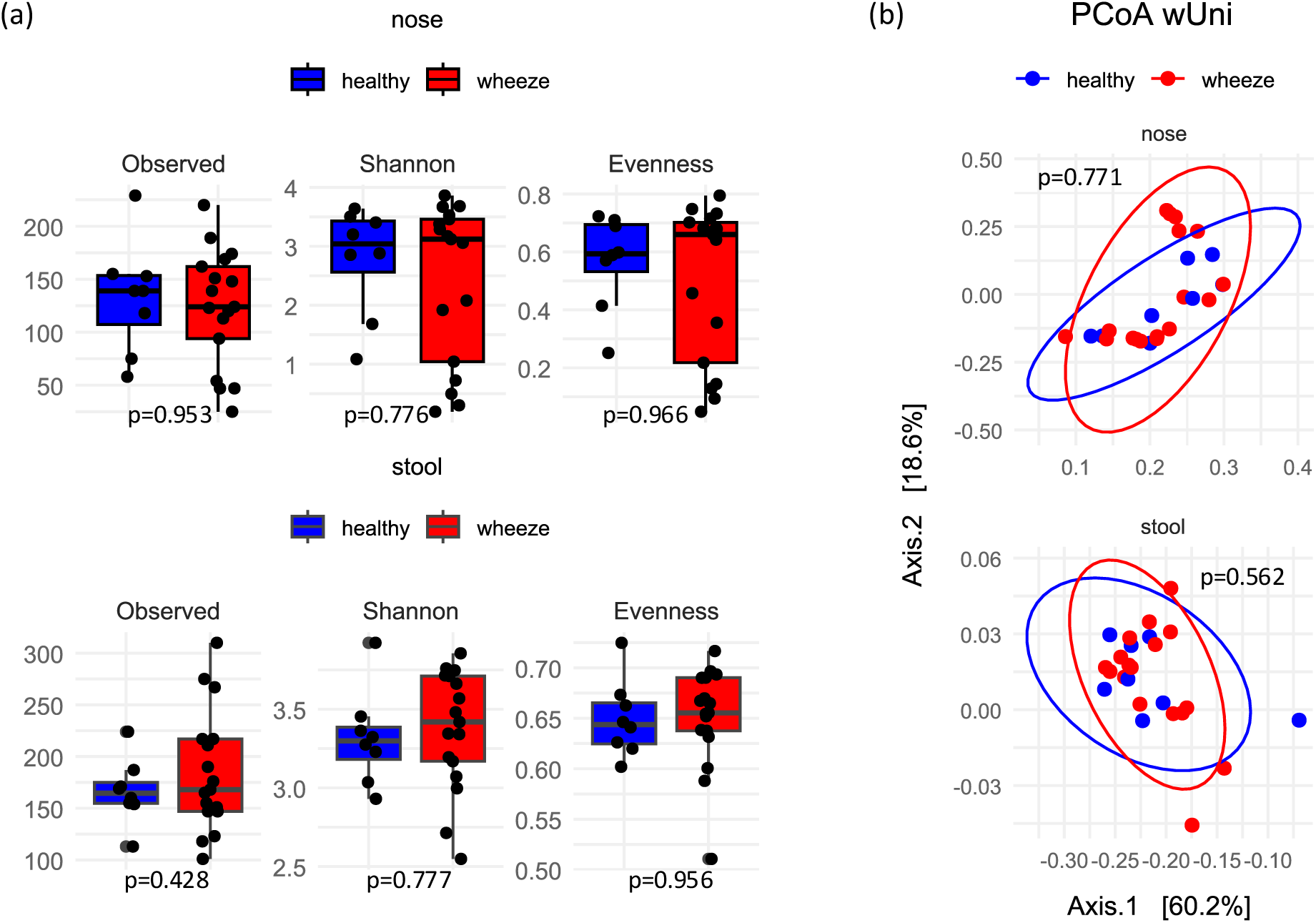
Microbial diversity and overall community composition in nose and stool of healthy (n=8) and wheezing (n=17) children. (a) Boxplots of alpha diversity measures (observed ASV, Shannon index and evenness). Group differences were assessed using generalized linear mixed-effects models with Benjamini–Hochberg adjustment. (b) Principal coordinates analysis (PCoA) of weighted UniFrac distances. Group differences were assessed using PERMANOVA with Benjamini–Hochberg correction.

**Figure S2.**
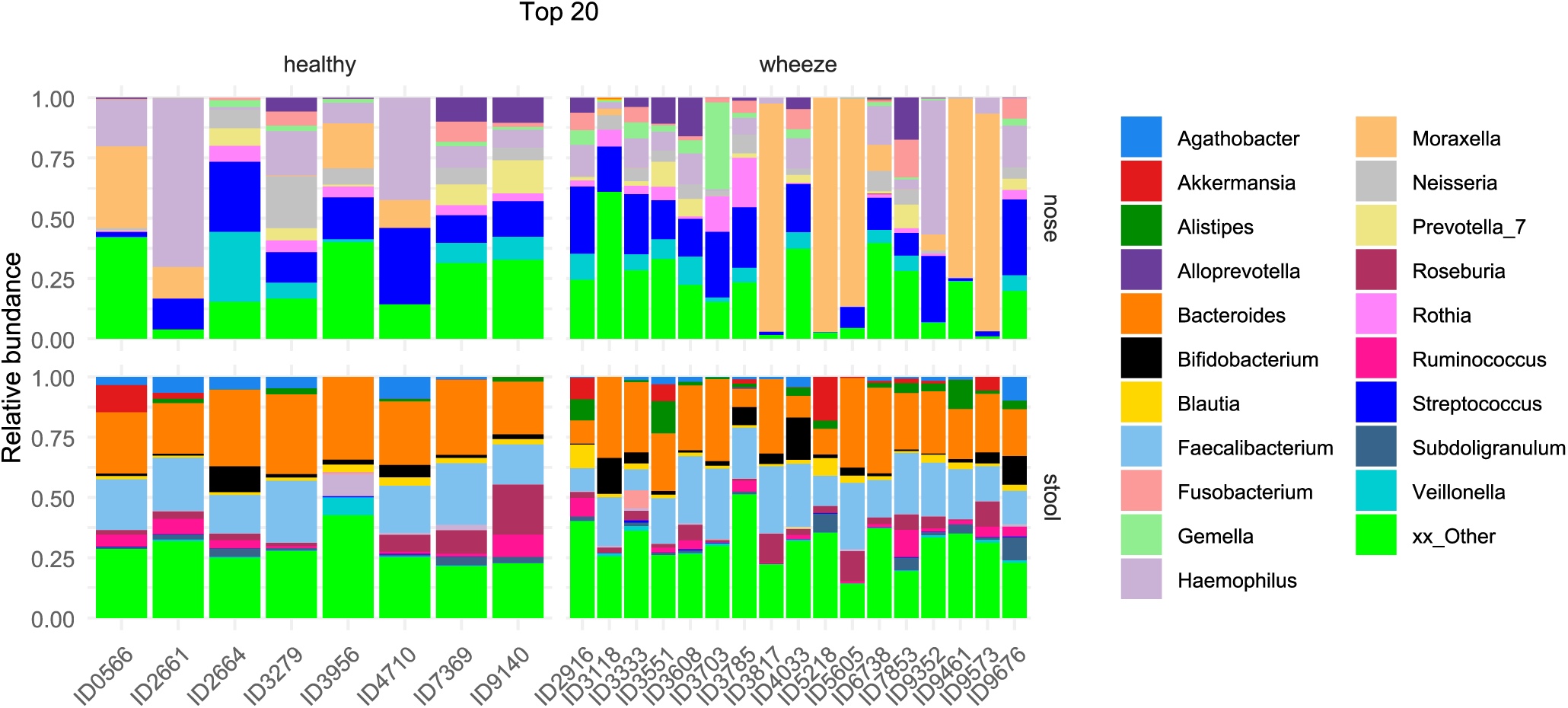
Relative abundance of the top 20 genera per subject.

**Figure S3.**
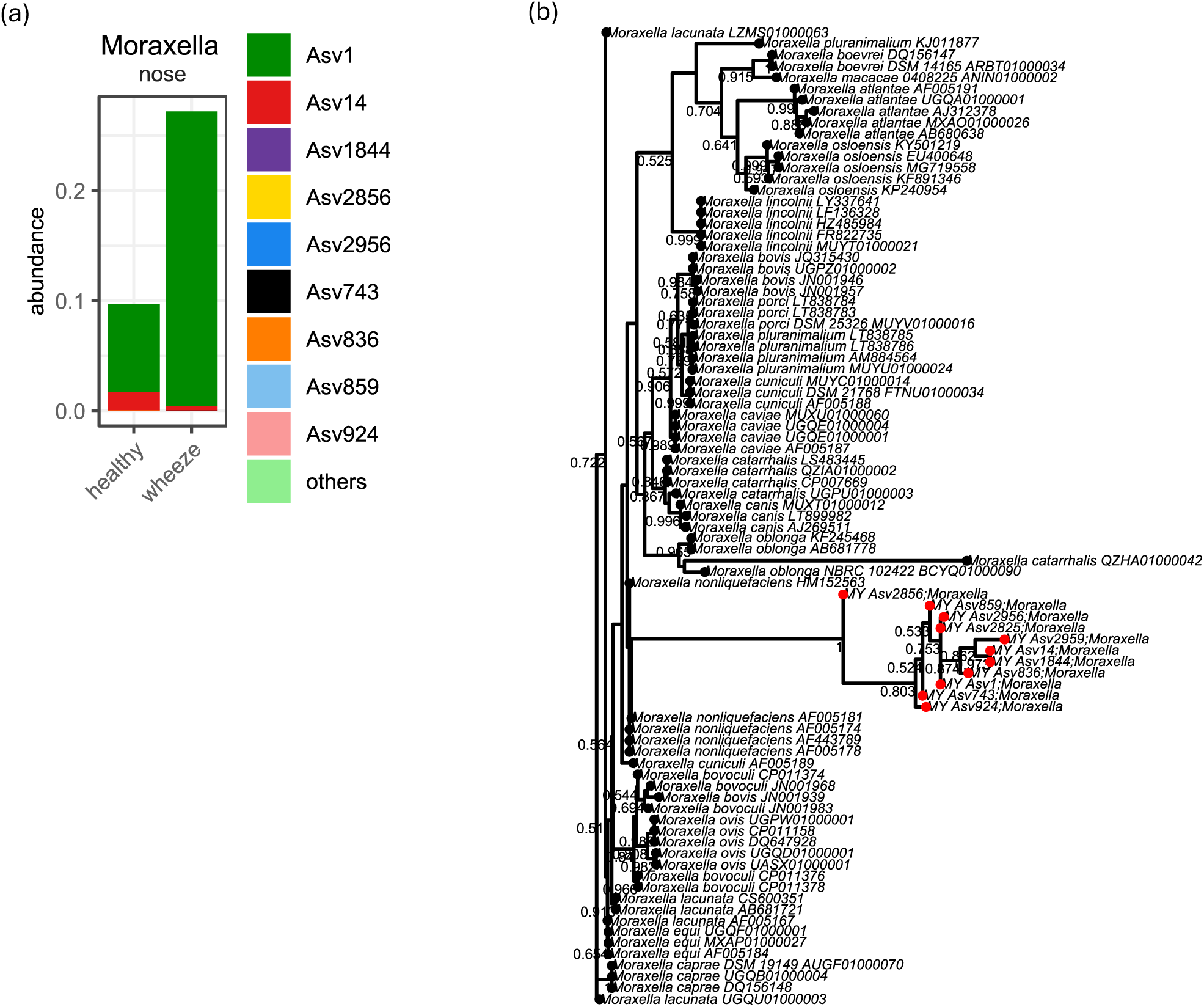
*Moraxella* ASV identified in the study cohort. (a) Relative abundance of *Moraxella* ASV in healthy and wheezing children. (b) Maximum-likelihood phylogenetic tree showing the placement of the detected *Moraxella* ASV (red dots) among reference sequences obtained from the SILVA database. Numbers at the nodes indicate bootstrap support values.

**Figure S4.**
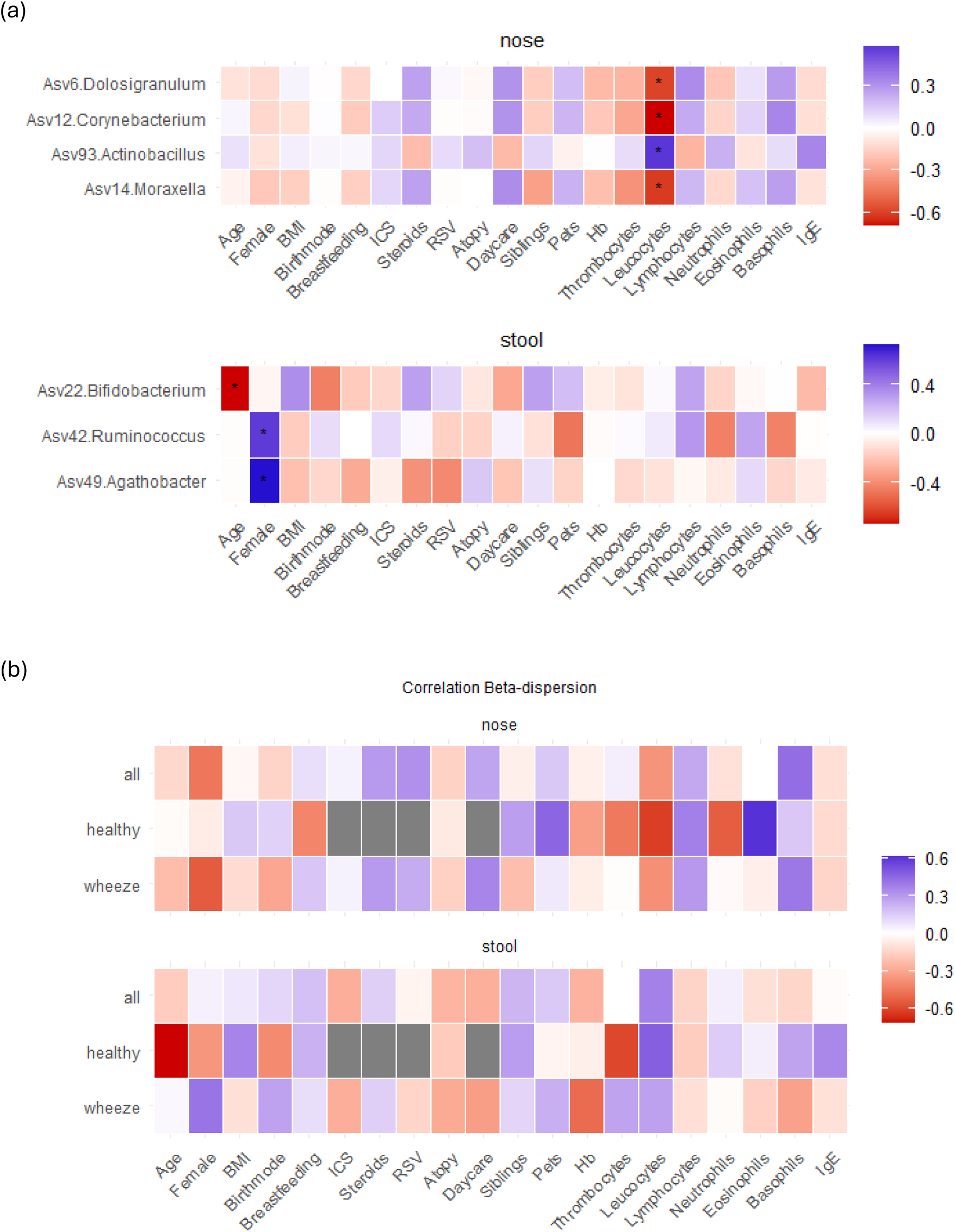
Associations between microbial community and demografic, lifestyle and immunological variables in nose and stool. (a) Spearman correlations between individual ASV and selected demographic, lifestyle and immunological variables in nasal and stool samples. Asterisks indicate statistically significant correlations after Benjamini-Hochberg adjustment (p<0.05), calculated using cor.test. (b) Pearson correlations between beta-dispersion and selected demographic, lifestyle and immunological factors in nasal and stool samples. Asterisks indicate statistically significant correlations after Benjamini-Hochberg adjustment (p<0.05), calculated using cor.test.

**Figure S5.**
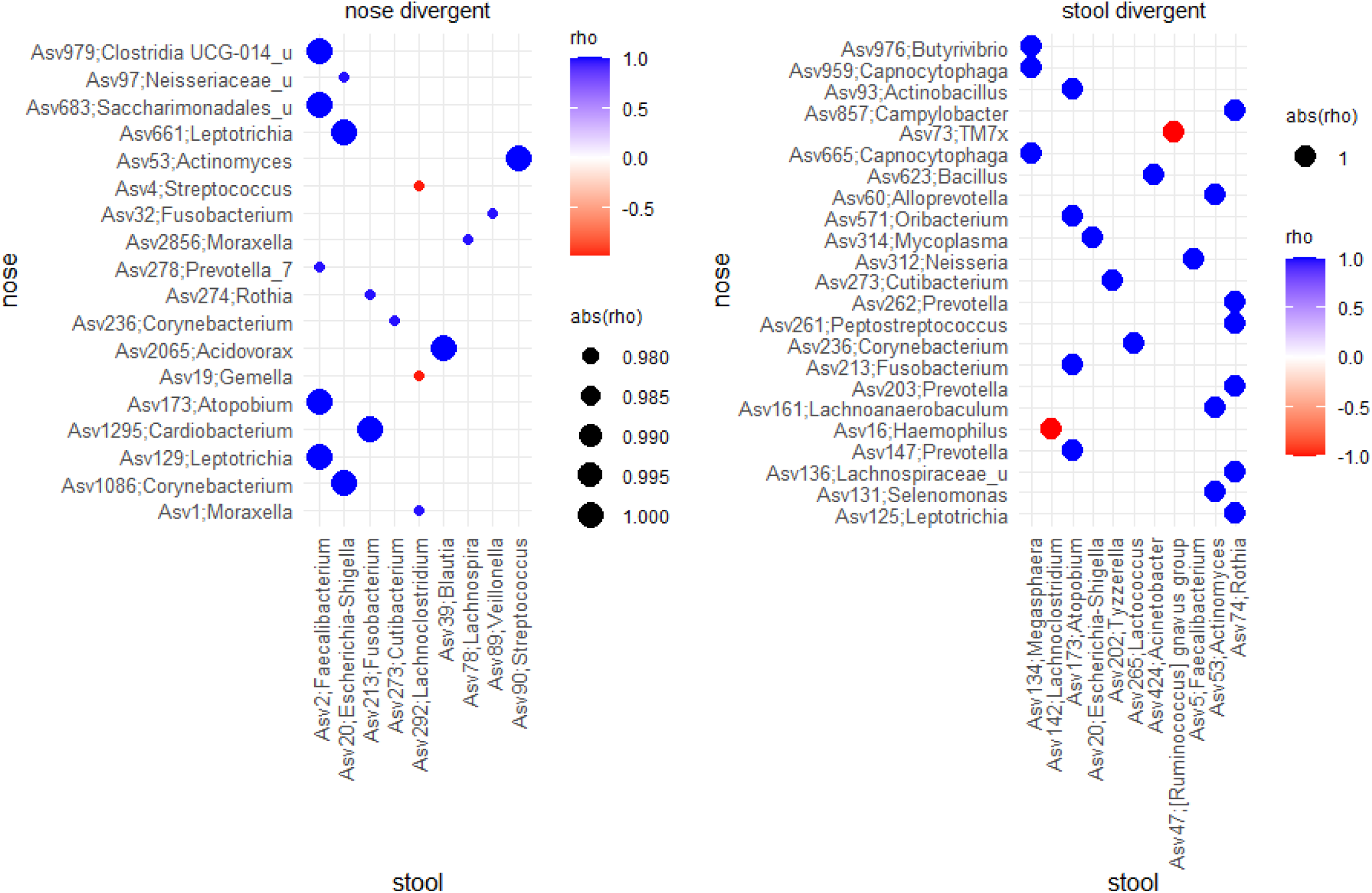
Pairwise cross-compartment correlations in community divergent subgroups. Spearman correlations were calculated between individual ASV in nasal (n=8) and stool (n=7) samples. Only statistically significant correlations after Benjamini–Hochberg adjustment (p<0.05) are shown.

**Supplementary Table S1.** Core ASV (≥80% prevalence within group) in nasal and stool samples, shown as binary presence/absence (1/0). Nose: 9 shared, 4 healthy-specific, 9 wheezer-specific ASV; stool: 30 shared, 11 healthy-specific, 13 wheezer-specific ASV.

| Taxa | nose |  | stool |  | Prev |
| --- | --- | --- | --- | --- | --- |
|  | Healthy | Wheeze | Healthy | Wheeze |  |
| Asv1;Moraxella | 1 | 0 | 0 | 0 | prev0.8 |
| Asv10;Veillonella | 0 | 1 | 1 | 1 | prev0.8 |
| Asv101;Clostridium sensu stricto 1 | 0 | 0 | 1 | 0 | prev0.8 |
| Asv105;Monoglobus | 0 | 0 | 1 | 1 | prev0.8 |
| Asv106;Clostridium sensu stricto 1 | 0 | 0 | 1 | 1 | prev0.8 |
| Asv110;Neisseria | 0 | 1 | 0 | 0 | prev0.8 |
| Asv111;Anaerostipes | 0 | 0 | 1 | 1 | prev0.8 |
| Asv114;Porphyromonas | 1 | 1 | 0 | 0 | prev0.8 |
| Asv116;Veillonella | 0 | 0 | 1 | 1 | prev0.8 |
| Asv13;Roseburia | 0 | 0 | 1 | 0 | prev0.8 |
| Asv132;Bifidobacterium | 0 | 0 | 0 | 1 | prev0.8 |
| Asv142;Lachnoclostridium | 0 | 0 | 1 | 1 | prev0.8 |
| Asv15;Bacteroides | 0 | 0 | 1 | 0 | prev0.8 |
| Asv152;Oscillibacter | 0 | 0 | 0 | 1 | prev0.8 |
| Asv158;Intestinibacter | 0 | 0 | 1 | 1 | prev0.8 |
| Asv16;Haemophilus | 0 | 1 | 1 | 1 | prev0.8 |
| Asv166;Clostridium sensu stricto 1 | 0 | 0 | 1 | 1 | prev0.8 |
| Asv167;Romboutsia | 0 | 0 | 1 | 1 | prev0.8 |
| Asv17;Prevotella_7 | 0 | 1 | 0 | 0 | prev0.8 |
| Asv172;Anaerostipes | 0 | 0 | 0 | 1 | prev0.8 |
| Asv18;Dialister | 0 | 0 | 1 | 0 | prev0.8 |
| Asv180;Porphyromonas | 0 | 1 | 0 | 0 | prev0.8 |
| Asv19;Gemella | 1 | 1 | 0 | 0 | prev0.8 |
| Asv2;Faecalibacterium | 0 | 0 | 1 | 1 | prev0.8 |
| Asv20;Escherichia-Shigella | 0 | 0 | 1 | 1 | prev0.8 |
| Asv209;Actinomyces | 0 | 1 | 0 | 0 | prev0.8 |
| Asv215;Erysipelatoclostridium | 0 | 0 | 1 | 1 | prev0.8 |
| Asv22;Bifidobacterium | 0 | 0 | 1 | 0 | prev0.8 |
| Asv23;Roseburia | 0 | 0 | 1 | 1 | prev0.8 |
| Asv234;Ruminococcaceae_u | 0 | 0 | 1 | 1 | prev0.8 |
| Asv24;Alistipes | 0 | 0 | 0 | 1 | prev0.8 |
| Asv25;Streptococcus | 1 | 1 | 1 | 1 | prev0.8 |
| Asv250;Hungatella | 0 | 0 | 1 | 1 | prev0.8 |
| Asv252;Lautropia | 1 | 0 | 0 | 0 | prev0.8 |
| Asv257;[Clostridium] innocuum group | 0 | 0 | 1 | 1 | prev0.8 |
| Asv267;[Eubacterium] hallii group | 0 | 0 | 0 | 1 | prev0.8 |
| Asv268;UBA1819 | 0 | 0 | 0 | 1 | prev0.8 |
| Asv27;Bifidobacterium | 0 | 0 | 1 | 1 | prev0.8 |
| Asv274;Rothia | 1 | 0 | 0 | 0 | prev0.8 |
| Asv28;Faecalibacterium | 0 | 0 | 1 | 0 | prev0.8 |
| Asv287;Eggerthella | 0 | 0 | 1 | 1 | prev0.8 |
| Asv29;Alloprevotella | 1 | 1 | 0 | 0 | prev0.8 |
| Asv292;Lachnoclostridium | 0 | 0 | 0 | 1 | prev0.8 |
| Asv3;Bacteroides | 0 | 0 | 0 | 1 | prev0.8 |
| Asv311;Lachnospiraceae NK4A136 group | 0 | 0 | 1 | 1 | prev0.8 |
| Asv33;Agathobacter | 0 | 0 | 1 | 1 | prev0.8 |
| Asv37;Neisseria | 0 | 1 | 0 | 0 | prev0.8 |
| Asv39;Blautia | 0 | 0 | 1 | 1 | prev0.8 |
| Asv4;Streptococcus | 1 | 1 | 1 | 0 | prev0.8 |
| Asv41;Bacteroides | 0 | 0 | 1 | 0 | prev0.8 |
| Asv42;Ruminococcus | 0 | 0 | 1 | 1 | prev0.8 |
| Asv45;Ruminococcus | 0 | 0 | 0 | 1 | prev0.8 |
| Asv47;[Ruminococcus] gnavus group | 0 | 0 | 1 | 1 | prev0.8 |
| Asv49;Agathobacter | 0 | 0 | 1 | 1 | prev0.8 |
| Asv5;Faecalibacterium | 0 | 0 | 1 | 1 | prev0.8 |
| Asv506;Anaerotruncus | 0 | 0 | 1 | 0 | prev0.8 |
| Asv51;Granulicatella | 1 | 1 | 0 | 0 | prev0.8 |
| Asv53;Actinomyces | 1 | 0 | 0 | 0 | prev0.8 |
| Asv61;Granulicatella | 1 | 1 | 0 | 0 | prev0.8 |
| Asv62;Actinomyces | 0 | 1 | 0 | 0 | prev0.8 |
| Asv64;Streptococcus | 1 | 1 | 1 | 0 | prev0.8 |
| Asv657;Terrisporobacter | 0 | 0 | 1 | 0 | prev0.8 |
| Asv67;Blautia | 0 | 0 | 0 | 1 | prev0.8 |
| Asv71;Blautia | 0 | 0 | 0 | 1 | prev0.8 |
| Asv77;Flavonifractor | 0 | 0 | 1 | 1 | prev0.8 |
| Asv8;Haemophilus | 1 | 1 | 0 | 0 | prev0.8 |
| Asv85;Veillonella | 0 | 1 | 0 | 0 | prev0.8 |
| Asv86;Lachnospiraceae NK4A136 group | 0 | 0 | 1 | 1 | prev0.8 |
| Asv88;Fusicatenibacter | 0 | 0 | 1 | 1 | prev0.8 |
| Asv89;Veillonella | 0 | 0 | 0 | 1 | prev0.8 |
| Asv95;Lachnospiraceae UCG-004 | 0 | 0 | 0 | 1 | prev0.8 |

**Supplementary Table S2.** Summary of baseline characteristics of wheezing children further stratified by microbial community composition distances.

| Characteristic | Overall<br>N = 17 | nose |  |  | stool |  |  |
| --- | --- | --- | --- | --- | --- | --- | --- |
|  |  | healthy-like<br>N = 9 | divergent<br>N = 8 | p<br>value <sup>1</sup> | healthy-like<br>N = 10 | divergent<br>N = 7 | p<br>value <sup>1</sup> |
| Age (year) | 3.00 (1.90-3.40) | 3.00 (2.60-3.30) | 2.55 (1.50-3.70) | 0.772 | 3.00 (1.90-3.40) | 3.20 (1.50-3.70) | 0.807 |
| Female | 4 (24%) | 4 (44%) | 0 (0%) | 0.082 | 3 (30%) | 1 (14%) | 0.603 |
| BMI (kg/m<sup>3</sup>) | 16.98 (15.50-18.20) | 17.40 (15.50-18.20) | 16.67 (15.76-18.08) | 0.501 | 16.84 (15.30-18.20) | 17.90 (15.50-18.26) | 0.696 |
| Any sibling | 12 (71%) | 7 (78%) | 5 (63%) | 0.620 | 7 (70%) | 5 (71%) | >0.999 |
| Daycare | 15 (88%) | 7 (78%) | 8 (100%) | 0.471 | 9 (90%) | 6 (86%) | >0.999 |
| Pet in household (cat/dog) | 7 (41%) | 4 (44%) | 3 (38%) | >0.999 | 2 (20%) | 5 (71%) | 0.058 |
| Asthma or recurrent spastic/obstructive bronchitis - DD | 2 (13%) | 2 (22%) | 0 (0%) | 0.475 | 0 (0%) | 2 (33%) | 0.125 |
| Missing | 1 | 0 | 1 |  | 0 | 1 |  |
| Hayfever - DD | 1 (5.9%) | 1 (11%) | 0 (0%) | >0.999 | 1 (10%) | 0 (0%) | >0.999 |
| Eczema - DD | 2 (13%) | 1 (11%) | 1 (14%) | >0.999 | 1 (11%) | 1 (14%) | >0.999 |
| Missing | 1 | 0 | 1 |  | 1 | 0 |  |
| Food allergy - DD | 0 (0%) | 0 (0%) | 0 (0%) |  | 0 (0%) | 0 (0%) | -- <sup>2</sup> |
| Birthmode |  |  |  | 0.091 |  |  | 0.449 |
| Vaginal delivery | 8 (47%) | 5 (56%) | 3 (38%) |  | 6 (60%) | 2 (29%) |  |
| Primary cesarean section | 6 (35%) | 1 (11%) | 5 (63%) |  | 2 (20%) | 4 (57%) |  |
| Secondary cesarean section | 1 (5.9%) | 1 (11%) | 0 (0%) |  | 1 (10%) | 0 (0%) |  |
| Operative vaginal delivery | 2 (12%) | 2 (22%) | 0 (0%) |  | 1 (10%) | 1 (14%) |  |
| Any exclusive breastfeeding ever | 14 (82%) | 7 (78%) | 7 (88%) | >0.999 | 8 (80%) | 6 (86%) | >0.999 |
| Any hypoallergenic formula ever | 8 (47%) | 5 (56%) | 3 (38%) | 0.637 | 5 (50%) | 3 (43%) | >0.999 |
| Prior RSV infection ever | 7 (47%) | 3 (38%) | 4 (57%) | 0.619 | 3 (38%) | 4 (57%) | 0.619 |
| Missing | 2 | 1 | 1 |  | 2 | 0 |  |
| Antibiotics given for recurrent cough ever | 1 (33%) | 1 (50%) | 0 (0%) | >0.999 | 1 (33%) | NA | -- <sup>2</sup> |
| Missing | 14 | 7 | 7 |  | 7 | 7 |  |
| Systemic steroids ever | 9 (53%) | 4 (44%) | 5 (63%) | 0.637 | 4 (40%) | 5 (71%) | 0.335 |
| ICS ever | 13 (76%) | 6 (67%) | 7 (88%) | 0.576 | 8 (80%) | 5 (71%) | >0.999 |
| Hb (g/dl) | 11.90 (11.45-12.25) | 11.90 (11.50-12.25) | 11.75 (11.45-12.40) | 0.958 | 11.90 (11.50-12.30) | 11.90 (11.30-12.20) | 0.558 |
| Missing | 1 | 1 | 0 |  | 1 | 0 |  |
| Thrombocytes (10<sup>3</sup>/μl) | 403 (372-443) | 403 (374-430) | 418 (318-449) | 0.958 | 397 (370-443) | 408 (374-455) | 0.491 |
| Missing | 1 | 1 | 0 |  | 1 | 0 |  |
| Leucocytes (10<sup>3</sup>/μl) | 9.27 (8.24-11.65) | 10.80 (9.27-11.95) | 8.52 (7.94-9.91) | 0.074 | 9.40 (8.25-10.78) | 9.03 (8.23-12.20) | 0.874 |
| Missing | 1 | 1 | 0 |  | 1 | 0 |  |
| Neutrophils (%) | 42 (36-55) | 48 (36-55) | 39 (33-51) | 0.563 | 40 (37-47) | 55 (21-60) | 0.427 |
| Missing | 1 | 1 | 0 |  | 1 | 0 |  |
| Lymphocytes (%) | 40 (36-47) | 38 (33-42) | 47 (38-59) | 0.115 | 41 (39-48) | 38 (30-45) | 0.427 |
| Missing | 1 | 1 | 0 |  | 1 | 0 |  |
| Eosinophils (%) | 1.50 (0.78-2.40) | 1.14 (0.47-2.20) | 2.00 (1.00-3.90) | 0.295 | 2.00 (1.00-2.40) | 0.90 (0.50-2.00) | 0.375 |
| Missing | 2 | 1 | 1 |  | 1 | 1 |  |
| Basophils (%) | 0.55 (0.08-1.00) | 0.25 (0.04-0.75) | 0.85 (0.50-1.00) | 0.194 | 0.70 (0.08-1.00) | 0.50 (0.10-0.60) | 0.841 |
| Missing | 3 | 1 | 2 |  | 1 | 2 |  |
| IgE (IU/ml) | 19 (9-82) | 19 (9-93) | 26 (9-43) | 0.724 | 19 (9-43) | 37 (6-82) | 0.724 |
| Missing | 2 | 0 | 2 |  | 1 | 1 |  |
| Atopy (spec. IgE to any allergen ≥0.7 kU/L) | 5 (29%) | 3 (33%) | 2 (25%) | >0.999 | 4 (40%) | 1 (14%) | 0.338 |

Values are summarized as median and interquartile range (Q1-Q3) in brackets, or as total numbers per condition level and percentage in brackets.
DD = parental report of doctor's diagnosis
<sup>1</sup>Kruskal-Wallis rank sum test; Fisher's exact test
<sup>2</sup>p value not calculated due to insufficient variability (single-level data) and/or missing values

**Supplementary Table S3.** Alpha diversity indices and betadispersion of wheezing children further stratified by microbial community composition distances.

|  | nose |  |  | stool |  |  |
| --- | --- | --- | --- | --- | --- | --- |
|  | healthy-like | divergent | p | healthy-like | divergent | p |
| Observed ASV | 149.00 | 94.75 | <0.001 | 180.40 | 190.57 | 0.835 |
| Shannon index | 3.46 | 1.23 | <0.001 | 3.32 | 3.44 | 0.321 |
| Evenness | 0.69 | 0.28 | <0.001 | 0.64 | 0.66 | 0.236 |
| Betadispersion | 0.17 | 0.30 | <0.001 | 0.11 | 0.15 | 0.001 |

